# The ChAdOx1 vectored vaccine, AZD2816, induces strong immunogenicity against SARS-CoV-2 Beta (B.1.351) and other variants of concern in preclinical studies

**DOI:** 10.1101/2021.06.08.447308

**Authors:** Alexandra J Spencer, Susan Morris, Marta Ulaszewska, Claire Powers, Reshma Kailath, Cameron Bissett, Adam Truby, Nazia Thakur, Joseph Newman, Elizabeth R Allen, Indra Rudiansyah, Chang Liu, Wanwisa Dejnirattisai, Juthathip Mongkolsapaya, Hannah Davies, Francesca R Donnellan, David Pulido, Thomas P. Peacock, Wendy S. Barclay, Helen Bright, Kuishu Ren, Gavin Screaton, Patrick McTamney, Dalan Bailey, Sarah C Gilbert, Teresa Lambe

**Author notes:** Correspondence to: Alexandra J Spencer, The Jenner Institute, ORCRB, Roosevelt Drive, Oxford OX3 7DQ.

## Abstract

There is an ongoing global effort to design, manufacture, and clinically assess vaccines against SARS-CoV-2. Over the course of the ongoing pandemic a number of new SARS-CoV-2 virus isolates or variants of concern (VoC) have been identified containing mutations in key proteins. In this study we describe the generation and preclinical assessment of a ChAdOx1-vectored vaccine (AZD2816) which expresses the spike protein of the Beta VoC (B.1.351). We demonstrate that AZD2816 is immunogenic after a single dose. When AZD2816 is used as a booster dose in animals primed with a vaccine encoding the original spike protein (ChAdOx1 nCoV-19/ [AZD1222]), high titre binding and neutralising antibodies against Beta (B.1.351), Gamma (P.1) and Delta (B.1.617.2) are induced. In addition, a strong and polyfunctional T cell response was measured in these booster regimens. These data support the ongoing clinical development and testing of this new variant vaccine.

## Introduction

Since the first reports of infections caused by a novel coronavirus, there has been an unprecedented global effort to design, manufacture and test multiple vaccines against SARS-CoV-2. The majority of vaccines encode the full-length spike protein of SARS-CoV-2 and induce both T cell responses and neutralising antibodies to variable levels. COVID-19 vaccines are being deployed to varying degrees globally and real-world effectiveness data are demonstrating the positive impact vaccination is having on preventing COVID-related hospitalisation and death, irrespective of the level of neutralising antibodies induced against variants of concern (VoC)^1-4^ (https://assets.publishing.service.gov.uk/government/uploads/system/uploads/attachment_data/file/988619/Variants_of_Concern_VOC_Technical_Briefing_12_England.pdf).

Over the course of the pandemic a number of VoC have been identified, each containing multiple mutations within the viral genome. Variants with mutations in the spike protein, and in particular the receptor binding domain (RBD) which binds to the angiotensin converting enzyme-2 (ACE-2) receptor and facilitates viral cell entry, may escape vaccine-induced host immunity resulting in infection and disease. The Beta variant (B.1.351)^5^, first identified in October 2020, contains 10 changes across the spike protein with three amino acid changes in the RBD region (Figure 1). These changes in RBD are reported to increase binding between spike and ACE2, and also result in a reduced level of neutralising antibodies; however, T cell responses to peptides spanning the variant spike are still induced^6,7^.

**Figure 1:**
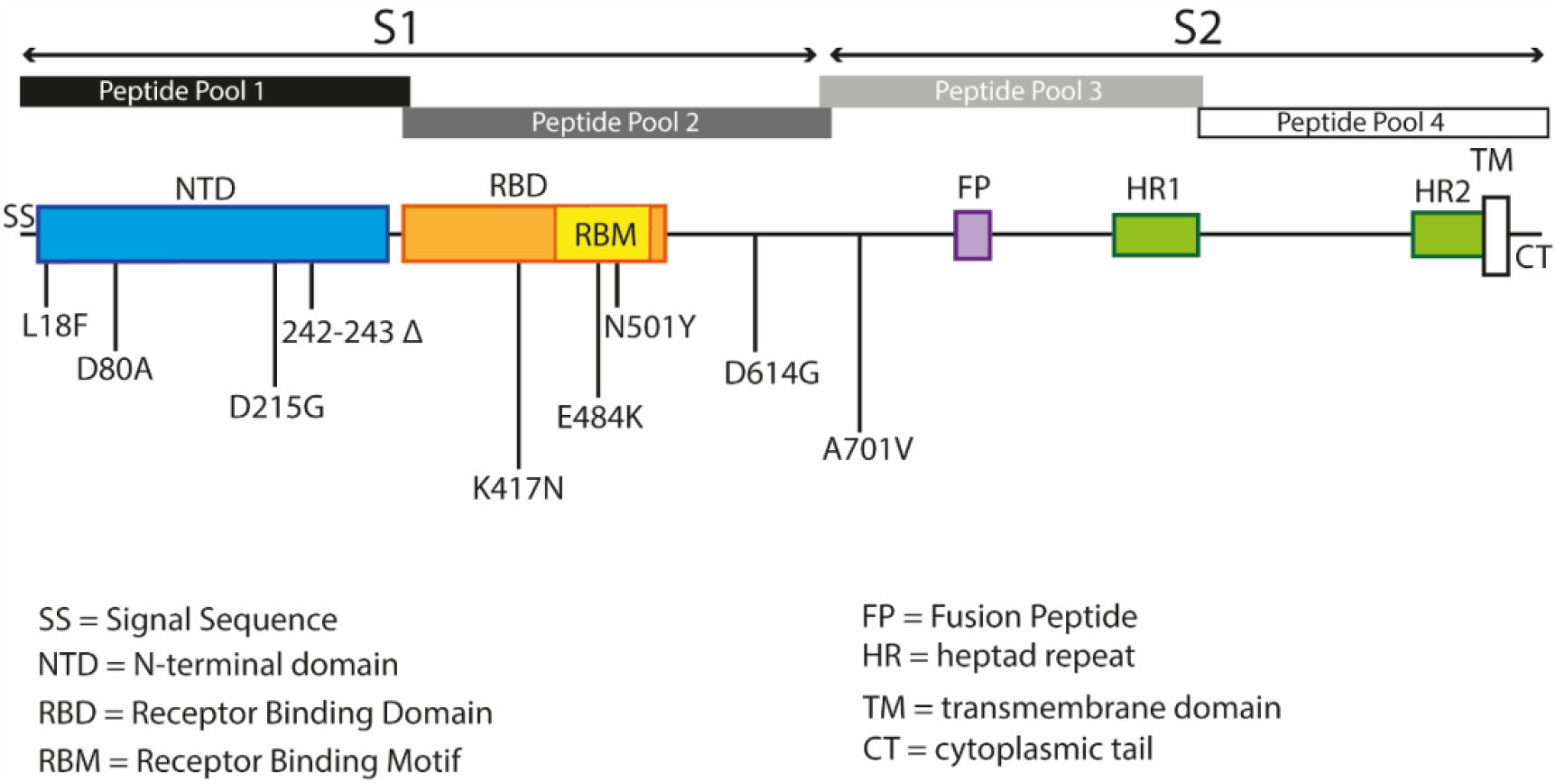
Schematic of SARS-CoV-2 spike protein and peptide pools used in studies. Schematic is a graphical representation of spike protein indicating location of the signal sequence (SS), N-terminal domain (NTD), receptor binding domain (RBD, receptor binding motif (RBM), fusion peptide (FP), heptad repeat (HR) regions, transmembrane domain (TM) and cytoplasmic tail (CT). Peptide pools used to stimulate splenocytes were sub-divided into 4 pools to cover the S1 and S2 regions of spike. Amino acid changes between original and Beta (B.1.351) variant virus and encoded in the AZD2816 vaccine construct are indicated. Triangle represents deletion of amino acids.

Platform vaccine technologies can be rapidly deployed to produce second generation SARS-CoV-2 vaccines targeting VoC. In this study we describe the generation and assessment of a ChAdOx1 expressing Beta spike protein (AZD2816) immunogenicity in mice. Importantly, a T cell immune response and both binding and neutralising antibodies against Beta were measured after a single dose vaccination with AZD2816. When AZD2816 was used as a booster dose in mice already primed with the original ChAdOx1 nCoV-19 (AZD1222) we observed strong antibody binding against both the original wild-type and Beta spike protein, with booster doses increasing the antibody response and neutralising ability against other variants. In a three-dose regimen using AZD2816 as a third dose, higher neutralising titres against VoC were induced. These data support the clinical testing of AZD2816 in prime-boost regimens after two doses of ChAdOx1 nCoV-19 vaccine as currently recommended.

## Results

### Single dose of AZD2816 vaccine induces cross-reactive immunity

Following reports of the SARS-CoV-2 beta variant (B.1.351), we generated a new ChAdOx1 vector expressing spike containing the key Beta (B.1.351) mutations (**Figure 1**) (AZD2816). Pre-fusion stabilisation has been reported to increase protein expression and improve immunogenicity of some viral glyco-proteins^8^, yet high level expression of pre-fusion spike protein on the surface of cells transfected with original ChAdOx1 nCov-19 (AZD1222), in which the antigen is not stabilised, has been reported ^9^, therefore the impact of spike stabilisation on immunogenicity could be vaccine platform dependant. ChAdOx1 expressing Beta spike protein containing 6 proline substitutions at aa 814, 889, 896, 939, 983 and 984 (hexa-pro)^8^, for pre-fusion stabilisation, was generated, and T cell and antibody responses following a single vaccination were compared with that of immunization with non-stabilised spike vaccination. Following a dose-range assessment of immune responses, in which a slightly lower T cell responses (**Figure S1B**), but no difference in antibody responses (**Figure S1A**) was observed between vaccines, further immunological assessment continued with the non-stabilised version (consistent with original ChAdOx1 nCoV-19 vaccine (AZD1222).

To compare the immunogenicity of ChAdOx1 nCoV-19 vaccines expressing different spike proteins, BALB/c mice were immunised with 10^8^ infectious units (iu) AZD1222 (ChAdOx1 nCoV-19), AZD2816 (ChAdOx1 nCoV-19 Beta) or with 10^8^ iu of each vaccine mixed together prior to immunisation (**Figure 2A**). Comparable levels of anti-spike antibodies were observed in all groups of vaccinated mice against both original spike and Beta (B.1.351) spike protein (**Figure 2B**). Mixing both vaccines together did not compromise the antibody response to either variant spike protein, nor was there a difference between total ELISA Units measured on day 9 or day 16 post-vaccination (**Figure 2B**). This rapid onset of a measurable antibody response suggests this vaccine is highly immunogenic. Neutralising antibodies, measured in a pseudotyped virus neutralisation assay, were detected against both the original and Beta spike (**Figure 2C**).

**Figure 2:**
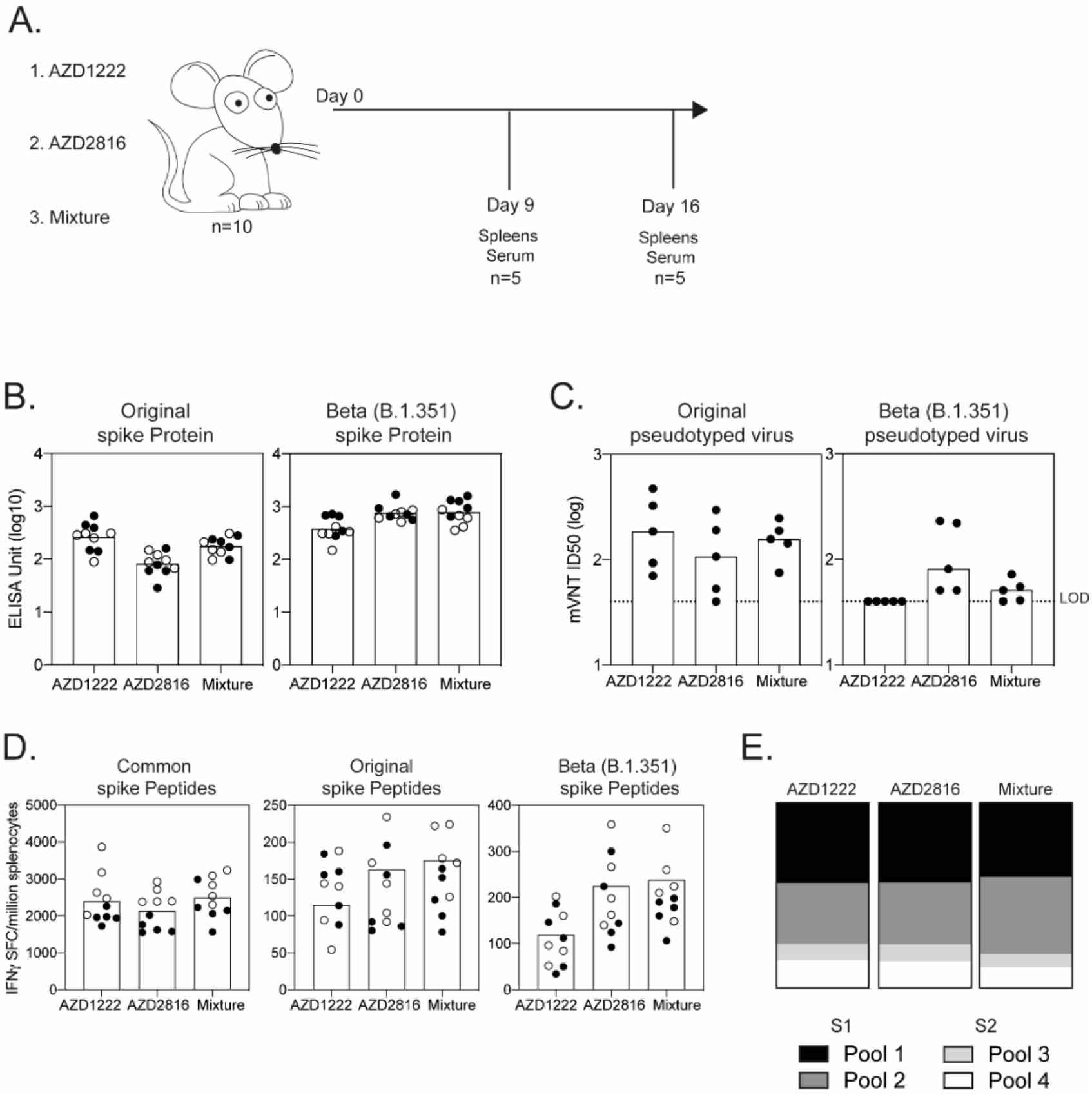
Immune response following a single dose of ChAdOx1 vaccines. **A**.) BALB/c mice (n=10) were vaccinated with 10^8^ iu of AZD1222 (ChAdOx1 nCoV-19), AZD2816 (ChAdOx1 nCoV-19 Beta) or 10^8^ iu of each vaccine mixed together. Mice were sacrificed 9 or 16 days later to measure antibody and T cell responses. **B**.) Spike-specific IgG levels measured in the serum of mice against original spike protein or Beta (B.1.351) spike protein. **C**.) Microneutralisation titres mVNT (ID50) measured in the serum of mice day 16 post vaccination, against pseudotyped virus expressing original spike or Beta (B.1.351) protein. Limit of detection (LOD) in the assay is defined as a titre of 40. **D**.) IFNγ secreting cells measured by ELISpot on day 9 or day 16, with splenocytes stimulated with pools of common peptides, original (WT) spike peptides or corresponding B.1.351 peptides covering the regions of difference between SARS-CoV-2 isolates. **E**.) Proportion of IFNγ secreting cells measured against spike common peptides, sub-divided into S1 (pool 1 and pool 2) or S2 (pool 3 or pool 4) regions of spike protein.

T cell responses were measured by IFNγ ELISpot with splenocytes stimulated with peptide pools containing peptides common to both spike antigens, or specific to peptides from the original or Beta strains (**Table S2**). Equivalent numbers of IFNγ producing cells after vaccination with AZD1222 or AZD2816 were detected at both timepoints measured (**Figure 2D**). T cell responses to the common peptides were dominant, with minimal responses observed against variant regions. Consistent with earlier studies ^10^, the T cell response was dominant towards the first 2 peptide pools corresponding to the S1 portion of the protein (**Figure 2E**) across all vaccine groups.

### Antibody responses are boosted by vaccination with variant vaccine AZD2816

Mice were immunised with one dose of AZD1222 prior to boosting 4 weeks later with AZD2816 and antibody responses compared a further 3 weeks later (**Figure 3A**). Total IgG levels, measured by ELISA, showed that a booster dose of AZD2816 increased the antibody titre against original spike and Beta spike (**Figure S2**). In addition, boosting AZD1222 primed mice with either AZD1222 or AZD2816 increased the binding and neutralising antibody titres against VoC, including original, Beta (B.1.351), Gamma (P.1), Delta (B.1.617.2), Epsilon (B.1.429) and Kappa (B.1.617.1) (**Figure 3B, S3 and Table 1**).

**Figure 3:**
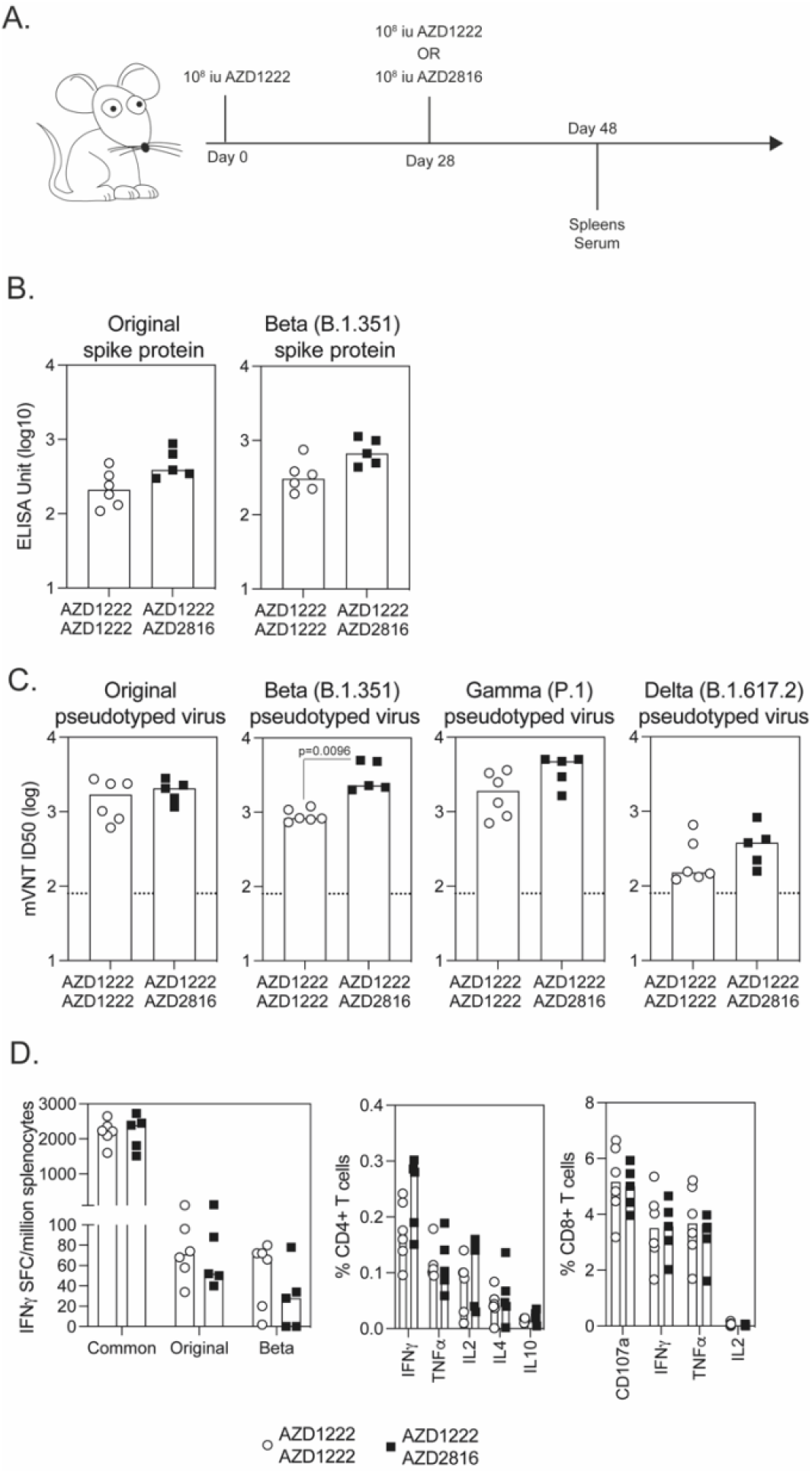
Immune response are boosted by immunisation with AZD2816. **A**.) BALB/c mice received one dose of 10^8^ iu of AZD1222 (ChAdOx1 nCoV-19) and were boosted with 10^8^ iu of AZD1222 or AZD2816 (ChAdOx1 nCoV-19 Beta). All mice were sacrificed a further 3 weeks later and antibody responses measured in the serum and T cell responses in the spleen of mice. **B**.) Graphs show the total IgG level measured by ELISA against original spike protein (WT) or B.1.351 spike protein. Data was log transformed and analysed with a two-way analysis of variance (repeated measure) and post-hoc positive test, no significance between groups (p<0.05) was observed. **C**.) Graphs show microneutralisation titres mVNT (ID50) measured against pseudotyped virus expressing original, Beta (B.1.351, Delta (B.1.617.2) or Gamma (P.1) spike protein. Limit of detection in the assay is defined as a titre of 80 (dotted line). Data was log transformed and analysed with a two-way analysis of variance (repeated measure) and post-hoc positive test, significance between groups (p<0.05) is indicated. **D**.) Graphs show IFNγ secreting cells measured by ELISpot, with splenocytes stimulated with pools of common, original (WT) or B.1.351 peptides or frequency of cytokine producing CD4^+^ (middle) or CD8^+^ T cells (right).

**Table 1:** Microneutralisation Titres.

| Prime | Boost | Boost | Time post last vaccine | Original wild-type spike |  | Beta (B.1.351) |  | Delta (B.1.617.2) |  | Gamma (P.1) |  |
| --- | --- | --- | --- | --- | --- | --- | --- | --- | --- | --- | --- |
|  |  |  |  | ID50 | ID80 | ID50 | ID80 | ID50 | ID80 | ID50 | ID80 |
| AZD1222 |  |  | 16 days | 186 (70 to 474) | 55 (43 to 297) | 40 | 40 | 40 | 40 (40 to 41) |  |  |
| AZD2816 |  |  | 16 days | 107 (40 to 297) | 40 (40 to 118) | 81 (51 to 231) | 55 (40 to 163) | 40 | 40 |  |  |
| AZD1222 & AZD2816 |  |  | 16 days | 157 (75 to 248) | 65 (40 to 93) | 51 (40 to 72) | 41 (40 to 51) | 40 | 40 |  |  |
| AZD1222 | AZD1222 |  | 20 days | 1691 (613 to 2750) | 486 (134 to 712) | 830 (729 to 1202) | 243 (129 to 485) | 151 (122 to 659) | 93 (80 to 342) | 2689 (1436 to 4861) | 722 (162 to 1457) |
| AZD1222 | AZD2816 |  | 20 days | 2058 (1159 to 2815) | 706 (477 to 926) | 2281 (2000 to 4984) | 585 (418 to 1371) | 379 (158 to 827) | 223 (85 to 454) | 3507 (3105 to 5100) | 998 (714 to 2049) |
| AZD1222 | AZD1222 | AZD1222 | 20 days | 4032 (2385 to 4559) | 1478 (507 to 2222) | 1609 (910 to 4519) | 413 (240 to 1617) | 721 (232 to 1274) | 375 (123 to 745) | 1896 (703 to 3610) | 1017 (406 to 2322) |
| AZD1222 | AZD1222 | AZD2816 | 20 days | 3704 (3462 to 4775) | 2022 (1135 to 3949) | 4392 (2304 to 4737) | 1864 (767 to 2844) | 1699 (547 to 4026) | 615 (200 to 1816) | 4755 (1637 to 5063) | 3236 (1294 to 4968) |

Functional ability of antibodies to neutralise pseudotyped virus expressing original spike, Beta (B.1.351), Delta (B.1.617.2) or Gamma (P.1) spike protein was measured in the serum of vaccinated mice. Pseudotyped virus neutralization titres are expressed as the reciprocal of the serum dilution that inhibited luciferase expression by 50% (ID50) or 80% (ID80). Table shows the median (min to max) per group.

Neutralising antibodies were observed in all animals following two doses of ChAdOx1 (either AZD1222/AZD1222 or AZD1222/AZD2816), with a statistically significant increase in neutralising antibodies against Beta pseudotyped virus observed in animals boosted with AZD2816 (**Figure 3C**).

Administering a booster vaccination after an initial dose of AZD1222 did not augment the T cell response, as has been reported before^11^. IFNγ ELISpot responses after a booster shot of either AZD1222 or AZD2816 were equivalent (**Figure 3D**), with T cell responses dominated by the response to common peptides (**Figure 3D**) and minimal responses detected against original or beta peptides. Priming with AZD1222 did not impact the polyfunctionality of the T cell response as the pattern of cytokine production from CD4^+^ or CD8^+^ T cells was similar in animals boosted with either AZD1222 or AZD2816 (**Figure 3D**).

### A third dose can further enhance antibody levels induced by two doses of AZD1222

AZD1222 has been authorised for use in a 2-dose vaccination regimen. We sought to determine if the immune response to a dose regimen could be enhanced with a booster dose of variant vaccine. A third dose of vaccine was administered to BALB/c mice which had previously received two doses of AZD1222 4 weeks apart and were then boosted a further 4 weeks later with 10^8^ iu of AZD2816 or AZD1222 (**Figure 4A**). An increase in spike-specific IgG was observed 3 weeks after the third dose of ChAdOx1, inducing higher spike-specific IgG compared with 2 doses (2.43 vs 2.87 Log10 spike-IgG titers **Figure 3A** and **Figure 4A**), regardless of the booster vaccine and against all variants of spike protein (**Figure 4B** and **S3B**).

**Figure 4:**
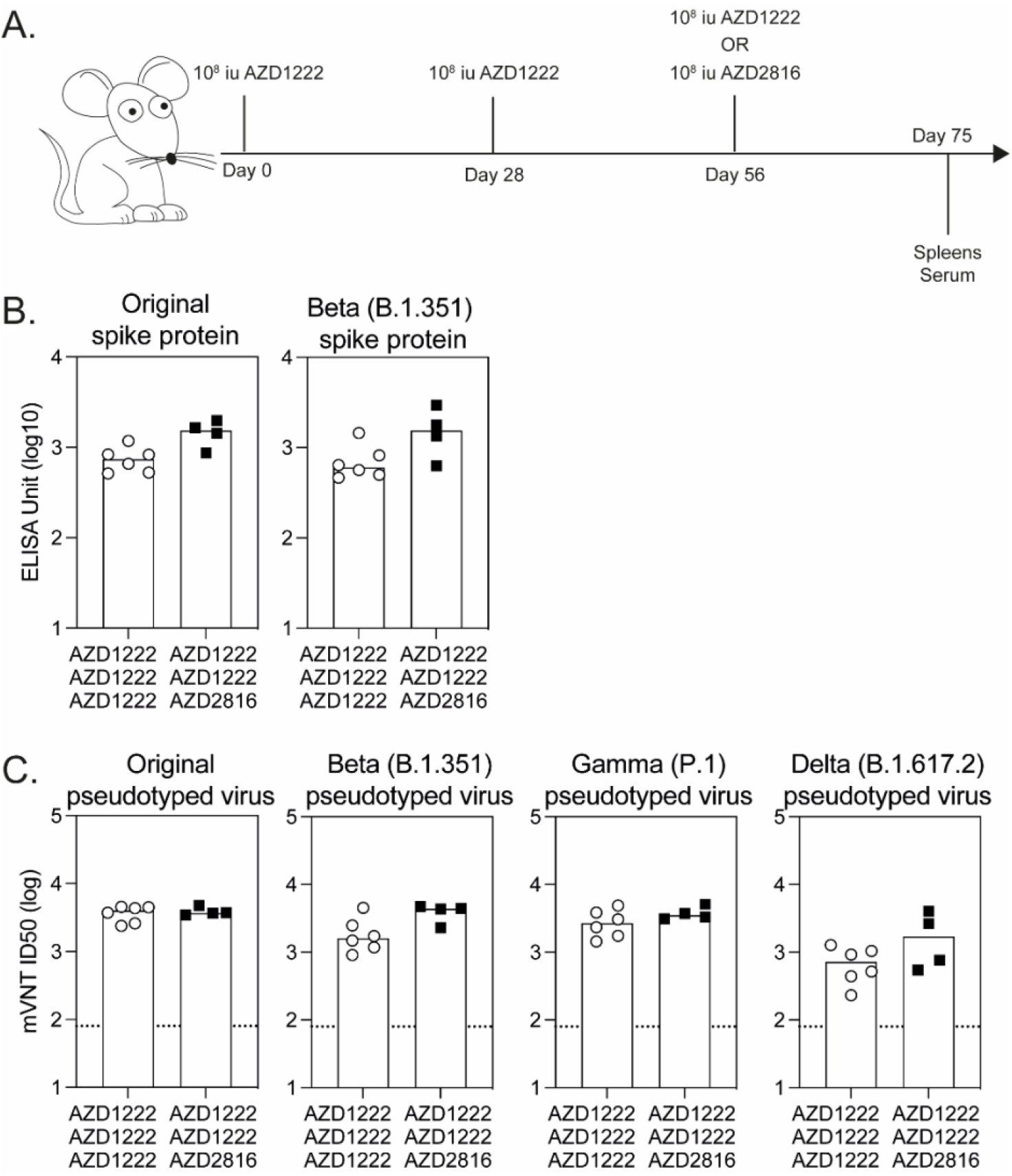
Immune response are boosted by immunisation with AZD2816. **A**.) BALB/c mice received two doses of 10^8^ iu of AZD1222 (ChAdOx1 nCoV-19) 4 weeks apart and were boosted with 10^8^ iu of AZD1222 or AZD2816 (ChAdOx1 nCoV-19 B.1.351). All mice were sacrificed a further 3 weeks later and antibody responses measured in the serum and T cell responses in the spleen of mice. **B**.) Graphs show the total IgG level measured by ELISA against original spike protein (WT) or B.1.351 spike protein. Data was log transformed and analysed with a two-way analysis of variance (repeated measure) and post-hoc positive test, no significance between groups (p<0.05) was observed. **C**.) Graphs show microneutralisation titres mVNT (ID80) measured against pseudotyped virus expressing original (WT), B.1.351, B.1.617.2 or P.1 spike protein. Limit of detection in the assay is defined as a titre of 80 (dotted line). Data was log transformed and analysed with a two-way analysis of variance (repeated measure) and post-hoc positive test, no significance between groups (p<0.05) was observed.

Neutralising antibody responses were detected in all vaccine groups against wild-type, Beta, Delta and Gamma spike protein pseudotyped virus (**Figure 4C** and **Table 1**) with significantly higher levels compared to 2 doses (3.19 vs 3.60 log10 mVNT ID50 **Figure 3C** and **Figure 4C Table 1**), although no significant differences between boosting with AZD1222 or AZD2816 were observed.

Although a booster dose with AZD2816 did not further increase the frequency of antigen specific T cells (**Figure 5**), the breadth of the T cell immune response remained consistent (**Figure 5**). Most T cells were specific to common SARS-CoV-2 spike peptides with minimal reactivity against peptides from either original spike or Beta spike (**Figure 5A**) as observed after a single dose of vaccine (**Figure 2**). Most importantly, a third immunization with AZD2816 did not alter T cell responses with CD4^+^ T cells shown to produce primarily IFNγ (**Figure 5B left**), and no significant difference in the proportion or number of T effector (Teff), T effector memory (Tem) or T central memory (Tcm) CD4^+^ T cells observed (**Figure 5B right**). Consistent with previous data in mice ^10^, the anti-spike cell-mediated response was predominantly CD8^+^ T cells, with a high frequency of CD8^+^ T cells producing IFNγ and TNFα observed in mice boosted with either AZD1222 or AZD2816 (**Figure 5C left**). The response was dominated by Teff and Tem CD8^+^ T cells and was similar between regimens involving a third booster of either AZD1222 or AZD2816 (**Figure 5C right**).

**Figure 5:**
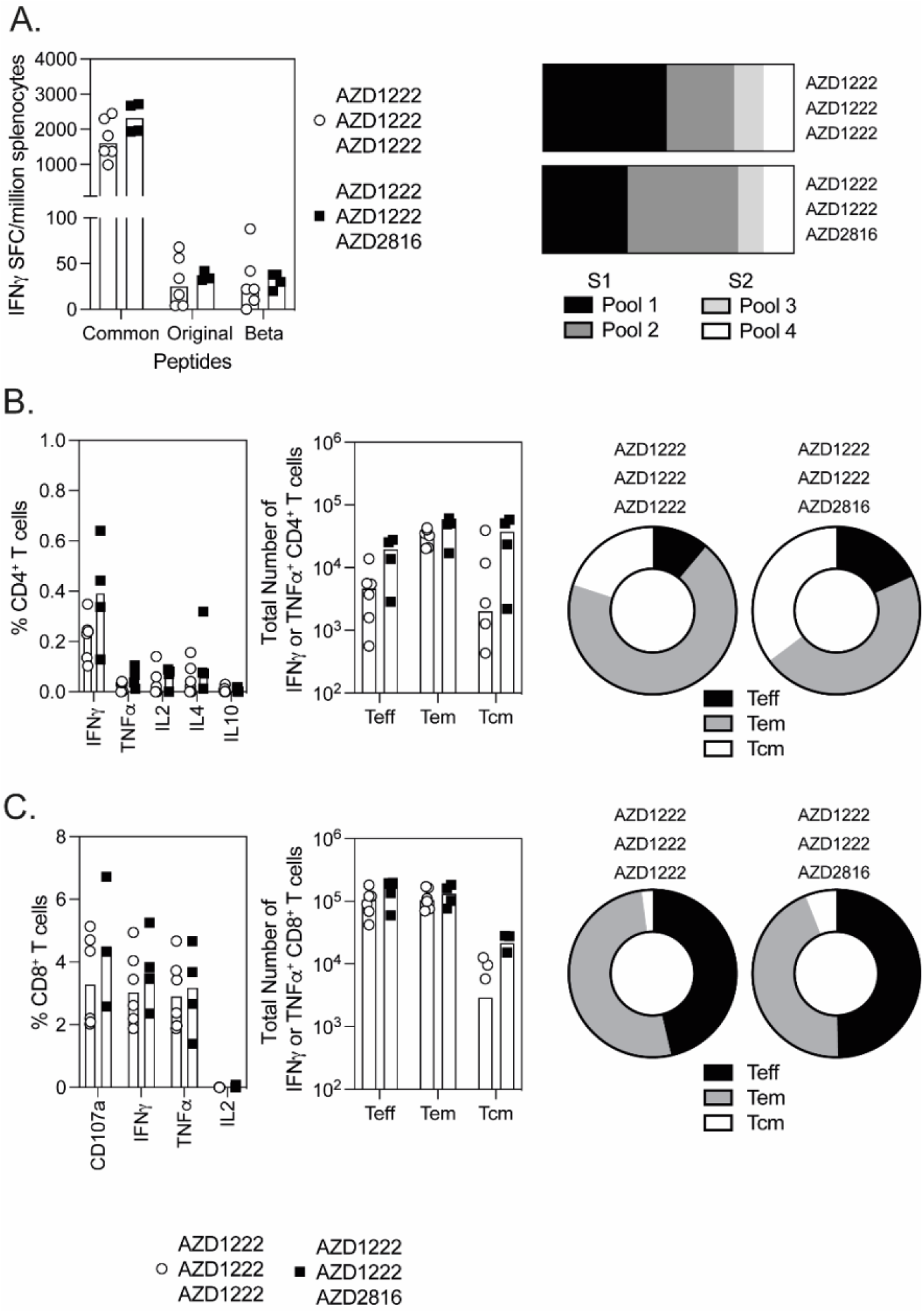
T cell responses following boost vaccination with AZD1222 or AZD2816. In the same experiment as described in Figure 4, T cell responses in the spleen of mice 3 weeks after the final vaccination. **A**.) Graphs show IFNγ secreting cells measured by ELISpot, with splenocytes stimulated with pools of common, original (WT) or B.1.351 peptides. Bar graph represents the proportion of IFNγ secreting cells measured against spike common peptides, sub-divided into S1 (pool 1 and pool 2) or S2 (pool 3 or pool 4) regions of spike protein. **B**.) Graphs show the frequency of cytokine producing CD4^+^, total number (left) or proportion (right) of IFNγ^+^ or TNFα^+^ CD4^+^ T cells of a T effector (Tem), T effector memory (Tem) or T central memory cells (Tcm) phenotype, bars represent the median response per group. **C**.) Graphs show the frequency of cytokine producing CD8^+^, total number (left) or proportion (right) of IFNγ^+^ or TNFα^+^ CD8^+^ T cells of a T effector (Tem), T effector memory (Tem) or T central memory cells (Tcm) phenotype, bars represent the median response per group.

Overall, the data shows that a booster dose with AZD2816 can further enhance antibody responses against the SARS-CoV-2 Beta VoC (B.1.351) and provide cross-reactivity against other spike variants, while maintaining robust and polyfunctional T cell responses.

## Discussion

A number of vaccine technologies, including viral vectors, allowed rapid production of vaccines against SARS-CoV-2 in early 2020 and can be rapidly deployed to generate new variant vaccines targeting the spike protein from VoCs. In this study we generated AZD2816, a new ChAdOx1 nCoV-19 vaccine encoding the Beta (B.1.351) spike protein and assessed the immunogenicity in mice post vaccination. We show that the use of proline stabilisation motif in the spike protein did not increase binding antibody titres after vaccination but was associated with a small reduction in T-cell responses (**Figure S1**). ChAdOx1 expressing non-stabilised Beta (B.1.351) spike was therefore selected for further development. This strategy was consistent with the original ChAdOx1 nCoV-19/AZD1222 vaccine design.

The Beta VoC (B.1.351), first identified in South Africa, contains several mutations across the S1 portion of spike protein. In particular, three mutations (K417N, E484K and N501Y) (**Figure 1**) involved in binding of spike to the ACE2 receptor have been shown to increase the avidity of the spike protein binding to ACE2, with sera from convalescent or vaccinated individuals showing reduced ability to neutralise this variant virus^6,12^. A number of common amino acid changes within the RBD and NTD region of the spike protein have been identified amongst SARS-CoV-2 variants (**Table S1**). The D614G change, identified in all VoC, increases virus infectivity ^13,14^ potentially through increased density of spike on the virion surface ^15^. The L452R change is present in Epsilon (B.1.429), Kappa (B.1.617.1) and Delta (B.1.617.2) and has been shown to reduce susceptibility to neutralising antibodies ^13^. The E484K change present in Beta (B.1.351) and Gamma (P.1) isolates and is thought to enhance binding affinity of RBD to ACE2 ^16,17^ and antibody evasion^18^. The N501Y change present in Beta (B.1.351), Alpha (B.1.1.7) and Gamma (P.1) variants alone, and does not appear to significantly impact neutralisation, but N501Y in combination with E484K and D614G can affect sera neutralisation titres ^19,20^. A high proportion of neutralising anti-spike antibodies bind to the RBD domain of spike ^21-23^, and there is concern that these cumulative changes are leading to the reduced ability of antibodies induced against WT SARS-CoV-2 to neutralise VoCs ^6,7,24,25^. Although, even with a reduced neutralising antibody titre against VoC, real world effectiveness data is demonstrating the ongoing positive impact these vaccines are having in preventing hospitalisation and death ^4^ (https://assets.publishing.service.gov.uk/government/uploads/system/uploads/attachment_data/file/988619/Variants_of_Concern_VOC_Technical_Briefing_12_England.pdf).

The initial exposure to a pathogen will prime a specific immune response and subsequent exposures are impacted by this pre-existing immunity including B cells and T cells specific to the original strain. This phenomenon, known as “original antigenic sin” or “antigenic imprinting” is commonly observed in the influenza field and may have a role to play in coronavirus biology^26-28^. As priming of the immune response against the original wild-type spike protein may impact the ability to switch specificity of the response to the Beta VoC (B.1.351), we measured antibody and T cell responses after one or two doses of the original ChAdOx1 nCoV-19 vaccine (AZD1222) followed by a single dose of AZD2816. While a single dose of either AZD1222 or AZD2816 induces rapid T cells and antibodies capable of binding and neutralising wild-type and Beta (B.1.351) spike protein, antibody responses were increased with a booster dose of either AZD1222 or AZD2816. Importantly, we saw no evidence that priming of the immune response was detrimental when mice received a booster dose of ChAdOx1 expressing Beta (B.1.351) protein.

Ongoing surveillance has identified Delta (B.1.617.2) as a VoC that has now spread rapidly around the world. Two dose vaccination with AZD1222 induced antibodies capable of neutralising Delta (B.1.617.2) and Gamma (P.1) in mice. However, it has been demonstrated in clinical settings that neutralising titres against VoC are significantly lower post vaccination with a number of approved vaccines^12^. Nonetheless, accruing real-world data is demonstrating the effectiveness of vaccination at preventing hospitalisation and death even in regions where VoC are circulating (https://assets.publishing.service.gov.uk/government/uploads/system/uploads/attachment_data/file/988619/Variants_of_Concern_VOC_Technical_Briefing_12_England.pdf) ^4,29^. These data suggest that while neutralising titres have been correlated with vaccine efficacy^30^, there are likely other immune mediators at play which can protect against severe disease, including T cells^31^. We demonstrate that high levels of T cells were observed after a third dose vaccination regimen with equivalent cytokines produced and populations of effector and memory T cells whether animals received a third vaccination with either AZD1222 or AZD2816. These responses were polyfunctional and had a predominantly effector memory T cell phenotype, which has been associated with rapid responses upon re-encounter with the virus.

The data presented herein demonstrates that vaccination with ChAdOx1 viral vectored vaccines targeting SARS CoV-2 (AZD1222/AZD2816) induces high titre cross-reactive antibodies capable of neutralising a number of SARS-CoV-2 VoCs including: Beta (B.1.351), Gamma (P.1) and Delta (B.1.617.2). Most importantly, T cells responses are maintained and neutralising antibody titres against VoC can be further enhanced by a booster dose of vaccine expressing the spike protein from Beta (B.1.351). These data support clinical assessment of AZD2816 as a booster dose in ongoing clinical trials.

## Methods

### Vector Construction

AZD2816 vaccine was constructed by methods as previously described ^32^. In brief, the B.1.351 glycoprotein (S) gene ^5^ was codon-optimized for expression in human cell lines and synthesized with the tissue plasminogen activator (tPA) leader sequence at the 5′ end by GeneArt Gene Synthesis (Thermo Fisher Scientific). The S gene was inserted into the Gateway® recombination cassette of the shuttle plasmid containing a human cytomegalovirus major immediate early promoter (IE CMV), which includes intron A and two tetracycline operator 2 sites, and the bovine growth hormone polyadenylation signal. BACs containing the ChAdOx1 SARS-CoV-2 Beta (B.1.351) Spike protein were prepared by Gateway® recombination between the ChAdOx1 destination DNA BAC vector ^33^ and the shuttle plasmids containing the SARS CoV-2 S gene expression cassettes using standard protocols resulting in the insertion of the SARS-CoV-2 expression cassette at the E1 locus. The ChAdOx1 SARS CoV-2 S adenovirus genome was excised from the BAC using unique PmeI sites flanking the adenovirus genome sequence. ChAdOx1 SARS CoV-2 S viral vectors were rescued in T-REx™ cells (Invitrogen, Cat. R71007), a derivative of HEK293 cells which constitutively express the Tet repressor protein and prevent antigen expression during virus production. The resultant virus, ChAdOx1 nCov-19 B.1.351 (AZD2816), was purified by CsCl gradient ultracentrifugation as described previously. The titres were determined on T-REx™ cells using anti-hexon immunostaining assay based on the QuickTiter™ Adenovirus Titer Immunoassay kit (Cell Biolabs Inc).

### Ethics Statement

Mice were used in accordance with the UK Animals (Scientific Procedures) Act 1986 under project license number P9804B4F1 granted by the UK Home Office with approval from the local Animal Welfare and Ethical Review Board (AWERB) at the University of Oxford. Age matched animals were purchased from commercial suppliers as a batch for each experiment and randomly split into groups on arrival at our facility. Animals were group housed in IVCs under SPF conditions, with constant temperature (20-24°C) and humidity (45-65%) with lighting on a 13:11 light-dark cycle (7am to 8pm). For induction of short-term anaesthesia, animals were anaesthetised using vaporised IsoFlo^®^. All animals were humanely sacrificed at the end of each experiment by an approved Schedule 1 method.

### Animals and Immunizations

Inbred BALB/cOlaHsd (BALB/c) (Envigo) (n=5 to 7 mice per group), were immunized intramuscularly (i.m.) in the musculus tibialis with 10^8^ infectious units (iu) of ChAdOx1 vector. Mice were boosted with the relevant vaccine candidate 4 weeks later. All mice were sacrificed 3 weeks (or at a time indicated on figure legend) after the final vaccination with serum and spleens collected for analysis of humoral and cell-mediated immunity.

### Antigen specific IgG ELISA

MaxiSorp plates (Nunc) were coated with 250ng/well of full-length SARS-CoV-2 wild-type (original) spike (NC_045512), Beta (B.1.351) spike, Alpha (B.1.1.7) spike, Gamma (P.1) spike, Epsilon (B.1.429) spike and original wild-type spike sequence with a D to G amino acid substitution at position 614 (D614G) protein (Table S1) overnight at 4°C, prior to washing in PBS/Tween (0.05% v/v) and blocking with Blocker Casein in PBS (Thermo Fisher Scientific) for 1 hour at room temperature (RT). Standard positive serum (pool of mouse serum with high endpoint titre against original wild-type spike protein), individual mouse serum samples, negative and an internal control (diluted in casein) were incubated for 2 hrs at 21°C. Following washing, bound antibodies were detected by addition of a 1 in 5000 dilution of alkaline phosphatase (AP)-conjugated goat anti-mouse IgG (Sigma-Aldrich) for 1 hour at 21°C and addition of p-Nitrophenyl Phosphate, Disodium Salt substrate (Sigma-Aldrich). An arbitrary number of ELISA units (EU) were assigned to the reference pool and optical density values of each dilution were fitted to a 4-parameter logistic curve using SOFTmax PRO software. ELISA units were calculated for each sample using the optical density values of the sample and the parameters of the standard curve. All data was log-transformed for presentation and statistical analyses.

### Micro neutralisation test (mVNT) using lentiviral-based pseudotypes bearing the SARS-CoV-2 Spike

Spike-expressing plasmid constructs were generated using the QuikChange Lightning Multi Site-Directed Mutagenesis kit (Agilent) on a previously described Wuhan-hu-1 template^34^. Lentiviral-based SARS-CoV-2 pseudotyped viruses were generated in HEK293T cells incubated at 37 °C, 5% CO_2_ as previously described ^11^. Briefly, cells were seeded at a density of 7.5 × 10^5^ in 6 well dishes, before being transfected with plasmids as follows: 500 ng of SARS-CoV-2 spike (Original NC_045512, Beta B.1.351, Kappa B.1.617.1, Delta B.1.617.2, Gamma P.1) (Table S1), 600 ng p8.91 (encoding for HIV-1 gag-pol), 600 ng CSFLW (lentivirus backbone expressing a firefly luciferase reporter gene), in Opti-MEM (Gibco) along with 10 µL PEI (1 µg/mL) transfection reagent. A ‘no glycoprotein’ control was also set up using the pcDNA3.1 vector instead of the SARS-CoV-2 Spike expressing plasmid. The following day, the transfection mix was replaced with 3 mL DMEM with 10% FBS (DMEM-10%) and incubated for 48 and 72 hours, after which supernatants containing pseudotyped SARS-CoV-2 (SARS-CoV-2 pps) were harvested, pooled and centrifuged at 1,300 x g for 10 minutes at 4 °C to remove cellular debris. Target HEK293T cells, previously transfected with 500 ng of a human ACE2 expression plasmid (Addgene, Cambridge, MA, USA) were seeded at a density of 2 × 10^4^ in 100 µL DMEM-10% in a white flat-bottomed 96-well plate one day prior to harvesting SARS-CoV-2 pps. The following day, SARS-CoV-2 pps were titrated 10-fold on target cells, and the remainder stored at -80 °C. For mVNTs, sera was diluted 1 in 20 or 1 in 40 in serum-free media and 50 µL was added to a 96-well plate in triplicate and titrated 2-fold. A fixed-titre volume of SARS-CoV-2 pps was added at a dilution equivalent to 10^5^ to 10^6^ signal luciferase units in 50 µL DMEM-10% and incubated with sera for 1 hour at 37 °C, 5% CO2 (giving a final sera dilution of 1 in 40 or 1 in 80). Target cells expressing human ACE2 were then added at a density of 2 × 10^4^ in 100 µL and incubated at 37 °C, 5% CO2 for 72 hours. Firefly luciferase activity was then measured with BrightGlo luciferase reagent and a Glomax-Multi+ Detection System (Promega, Southampton, UK). Pseudotyped virus neutralisation titres were calculated by interpolating the point at which there was either 50% or 80% reduction in luciferase activity, relative to untreated controls (50% or 80% neutralisation, inhibitory dilution 50 or 80, ID50 or ID80).

### ELISpot and ICS staining

Spleen single cell suspension were prepared by passing cells through 70μM cell strainers and treatment with ammonium potassium chloride lysis solution prior to resuspension in complete media. Splenocytes were stimulated 15mer peptides (overlapping by 11) spanning the length of SARS-CoV-2 protein and tpa promoter, with peptide pools subdivided into common and variant peptide regions within the S1 and S2 region of spike (Figure 1A) (Table S2). For analysis of IFNγ production by ELISpot, splenocytes were stimulated with two pools of common S1 peptides (pools 1 and 2), two pools of common S2 peptides (pools 3 and 4) (final concentration of 2µg/mL) and pools of original or beta variant peptides on hydrophobic-PVDF ELISpot plates (Merck) coated with 5µg/mL anti-mouse IFNγ (AN18). After 18-20 hours of stimulation at 37°C, IFNγ spot forming cells (SFC) were detected by staining membranes with anti-mouse IFNγ biotin (1mg/mL) (R46A2) followed by streptavidin-Alkaline Phosphatase (1mg/mL) and development with AP conjugate substrate kit (BioRad, UK). Spots were enumerated using an AID ELISpot reader and software (AID).

For analysis of intracellular cytokine production, cells were stimulated at 37°C for 6 hours with 2μg/mL a pool of S1 (ELISpot pools 1 and 2) or S2 (ELISpot pools 3 and 4) total original spike peptides (Table S2), media or positive control cell stimulation cocktail (containing PMA-Ionomycin, BioLegend), together with 1μg/mL Golgi-plug (BD) and 2μl/mL CD107a-Alexa647 (Clone 1D4B). Following surface staining with CD3-A700 (Clone 17A2, 1 in 100), CD4-BUV496 (Clone GK1.5, 1 in 200), CD8-BUV395 (Clone 53-6.7, 1 in 200), CD11a-PECy7 (Clone H155-78, 1 in 200), CD44-BV780 (Clone IM7, 1 in 100), CD62L-BV711 (Clone MEL-14, 1 in 100), CD69-PECy7 (Clone H1.2F3, 1 in 100), CD103-APCCy7 (Clone 2E7, 1 in 100) and CD127-BV650 (Clone A7R34, 1 in 100) cells were fixed with 4% paraformaldehyde and stained intracellularly with IL2-PerCPCy5.5 (Clone JES6-5H4, 1 in 100), IL4-BV605 (Clone 11B11, 1 in 100), IL10-PE (Clone JES5-16E3, 1 in 100), IFNγ-e450 (Clone XMG1.2, 1 in 100) and TNFα-A488 (Clone MP6-XT22, 1 in 100) diluted in Perm-Wash buffer (BD). Sample acquisition was performed on a Fortessa (BD) and data analyzed in FlowJo V10 (TreeStar). An acquisition threshold was set at a minimum of 5000 events in the live CD3^+^ gate. Antigen specific T cells were identified by gating on LIVE/DEAD negative, size (FSC-A vs SSC), doublet negative (FSC-H vs FSC-A), CD3^+^, CD4^+^ or CD8^+^ cells and each individual cytokine. T cell subsets were gated within the population of “IFNγ^+^ or TNFα^+^” responses and are presented after subtraction of the background response detected in the corresponding media stimulated control sample for each mouse, and summing together the response detected to each pool of peptides. T effector (Teff) cells were defined as CD62L^low^ CD127^low^, T effector memory (Tem) cells defined as CD62L^low^ CD127^hi^ and T central memory (Tcm) cells defined as CD62L^hi^ CD127^hi^ (Figure S3). The total number of cells was calculated by multiplying the frequency of the background corrected population (expressed as a percentage of total lymphocytes) by the total number of lymphocytes counted in each individual spleen sample.

### Statistical analysis

All graphs and statistical analysis were performed using Prism v9 (Graphpad). For analysis of vaccination regimen against a single variable (eg IgG level), data was analysed with a one-way anova (Kruskal-Wallis) followed by post-hoc Dunn’s multiple comparison test. For analysis of vaccination regimen against multiple variables (eg each individual cytokine or T cell subset) the data was analysed with a two-way analysis of variance, where a significant difference was observed, a post-hoc analysis was performed to compare the overall effect of vaccination regimen. In graphs where a significant difference was observed between multiple vaccine groups, the highest p value is displayed on the graph. All data displayed on a logarithmic scale was log_10_ transformed prior to statistical analysis (ELISA Units, Neutralisation Titres, Total Cell Numbers).

## Data availability

The data that support the findings of this study are available within the article and its Supplementary Information files or are available from the corresponding author upon reasonable request. Source data are provided with this paper.

## Acknowledgments

The authors would like to thank the BMS staff for animal husbandry and A. Worth, J.Furze, M. Mykhaylyk and R. Evans for facilities support.

## Author Contributions

SM, RK, CP cloned and produced virus preparations; AJS, MU, AT, CB, ERA and IR performed animal procedures and/or sample processing; AJS, MU, NT, JN, CB performed experiments; AJS, NT, DA analyzed data; CL, WD, JM, HD, FRD, DP, TPP, WSB, HB, KR, GS, PM provided reagents; AJS, TL & SG designed the study. AJS & TL wrote the manuscript. All authors reviewed the final version of the manuscript.

---

**Table S1:** Sequence changes to SARS-CoV-2 spike protein.

|  | Original Sequence | Beta B.1.351 | Alpha B.1.1.7 | Gamma P.1 | Epsilon B.1.429 | D614G | Kappa B.1.617.1 | Delta B.1.617.2 |
| --- | --- | --- | --- | --- | --- | --- | --- | --- |
| **NTD** | S13 |  |  |  | I |  |  |  |
|  | L18 | F |  | F |  |  |  |  |
|  | L19 |  |  |  |  |  |  | R |
|  | T20 |  |  | N |  |  |  |  |
|  | P26 |  |  | S |  |  |  |  |
|  | H69-V70 |  | Δ |  |  |  |  |  |
|  | D80 | A |  |  |  |  |  |  |
|  | T95 |  |  |  |  |  | I |  |
|  | D138 |  |  | Y |  |  |  |  |
|  | G142 |  |  |  |  |  | D | D |
|  | Y144 |  | Δ |  |  |  |  |  |
|  | W152 |  |  |  | C |  |  |  |
|  | E154 |  |  |  |  |  | K |  |
|  | E156-F157 |  |  |  |  |  |  | Δ |
|  | R158 |  |  |  |  |  |  | G |
|  | R190 |  |  | S |  |  |  |  |
|  | D215 | G |  |  |  |  |  |  |
|  | L242-A243 | Δ |  |  |  |  |  |  |
| **RBD** | K417 | N |  | T |  |  |  |  |
|  | L452 |  |  |  | R |  | R | R |
|  | T478 |  |  |  |  |  |  | K |
|  | E484 | K |  | K |  |  | Q |  |
|  | N501 | Y | Y | Y |  |  |  |  |
| **Other** | A570 |  | D |  |  |  |  |  |
|  | D614 | G | G | G | G | G | G | G |
|  | H655 |  |  | Y |  |  |  |  |
|  | P681 |  | H |  |  |  | R | R |
|  | A701 | V |  |  |  |  |  |  |
|  | T716 |  | I |  |  |  |  |  |
|  | D950 |  |  |  |  |  |  | N |
|  | S982 |  | A |  |  |  |  |  |
|  | T1027 |  |  | I |  |  |  |  |
|  | Q1071 |  |  |  |  |  | H |  |
|  | D1118 |  | H |  |  |  |  |  |
|  | V1176 |  |  | F |  |  |  |  |

**Table S2:**
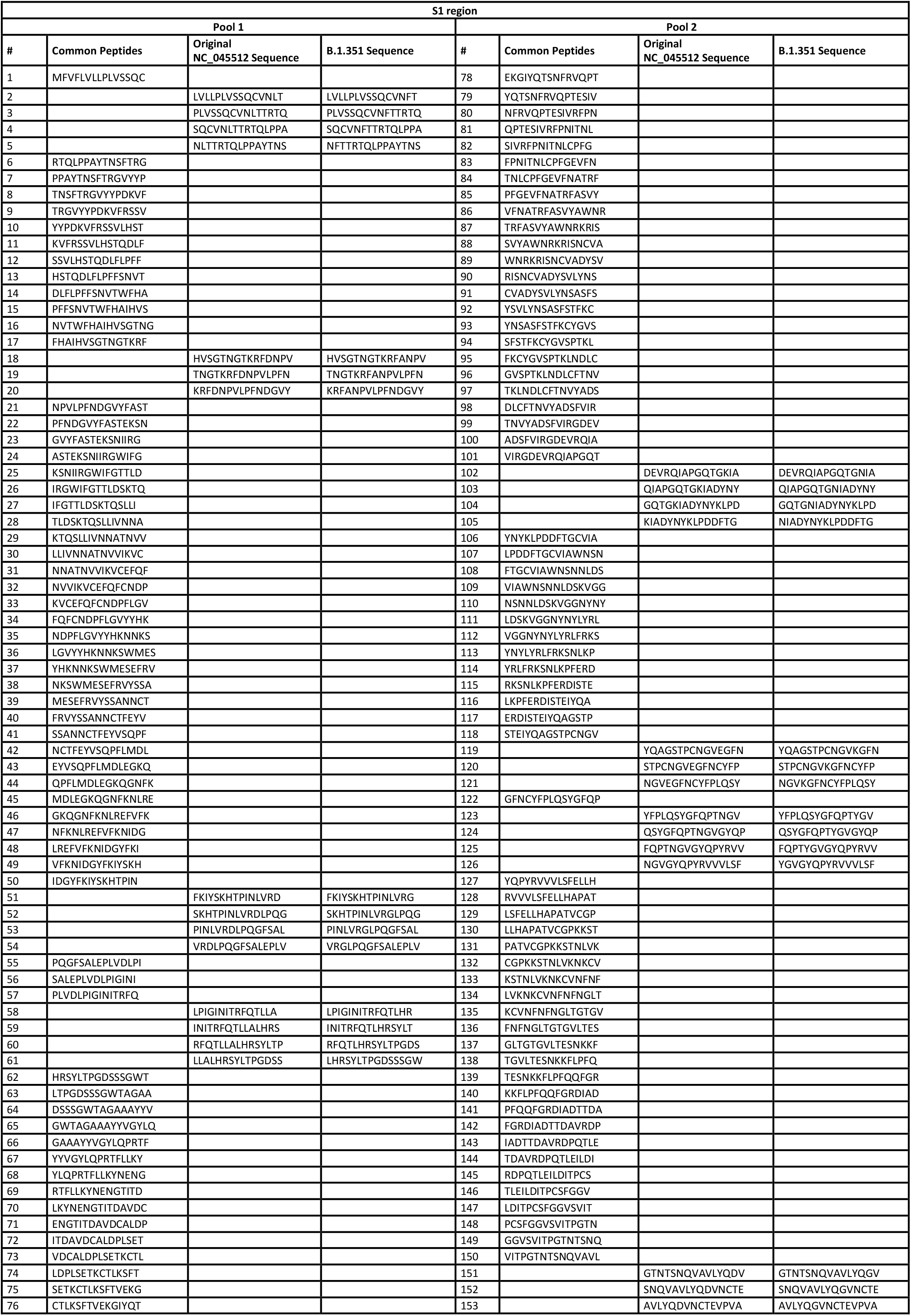
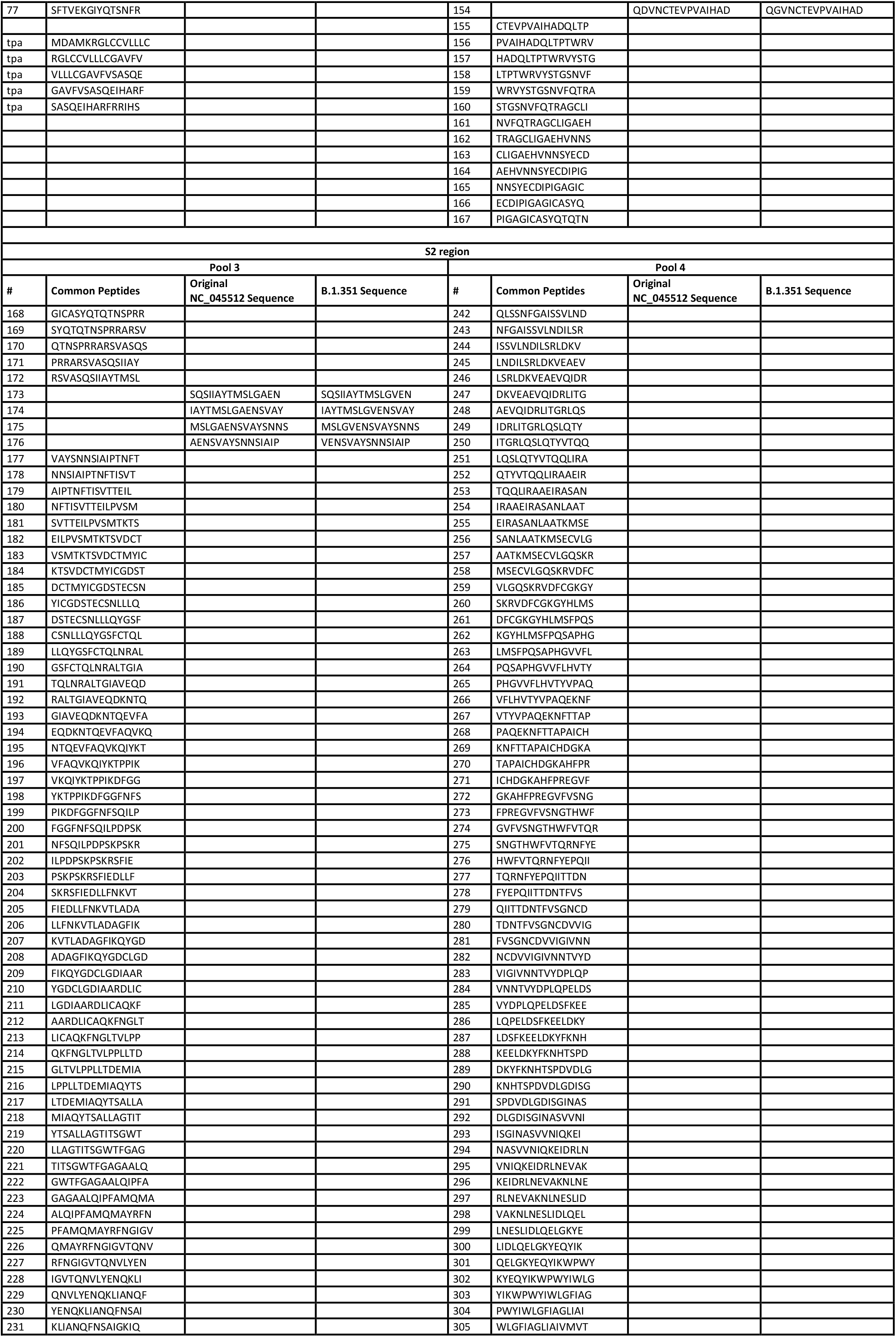
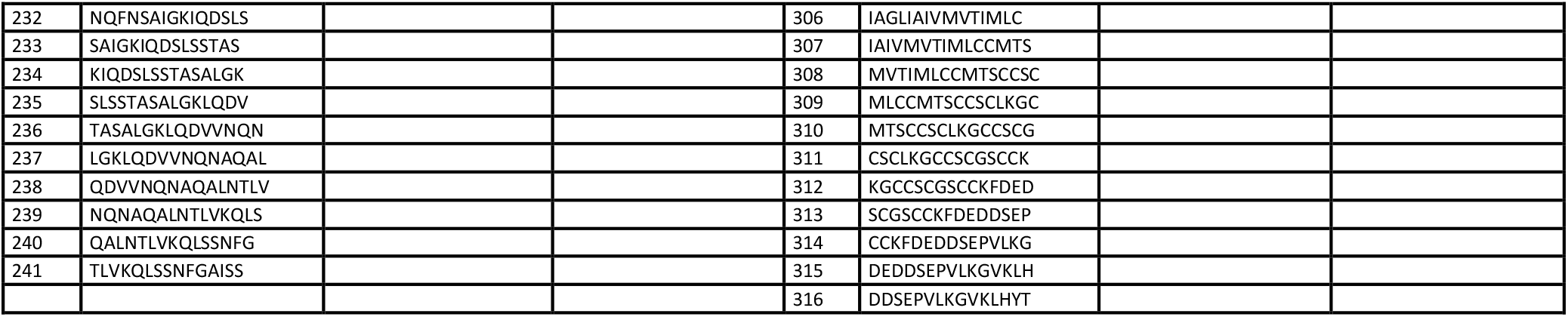
Overlapping SARS-CoV-2 spike peptide sequences.

**Figure S1:**
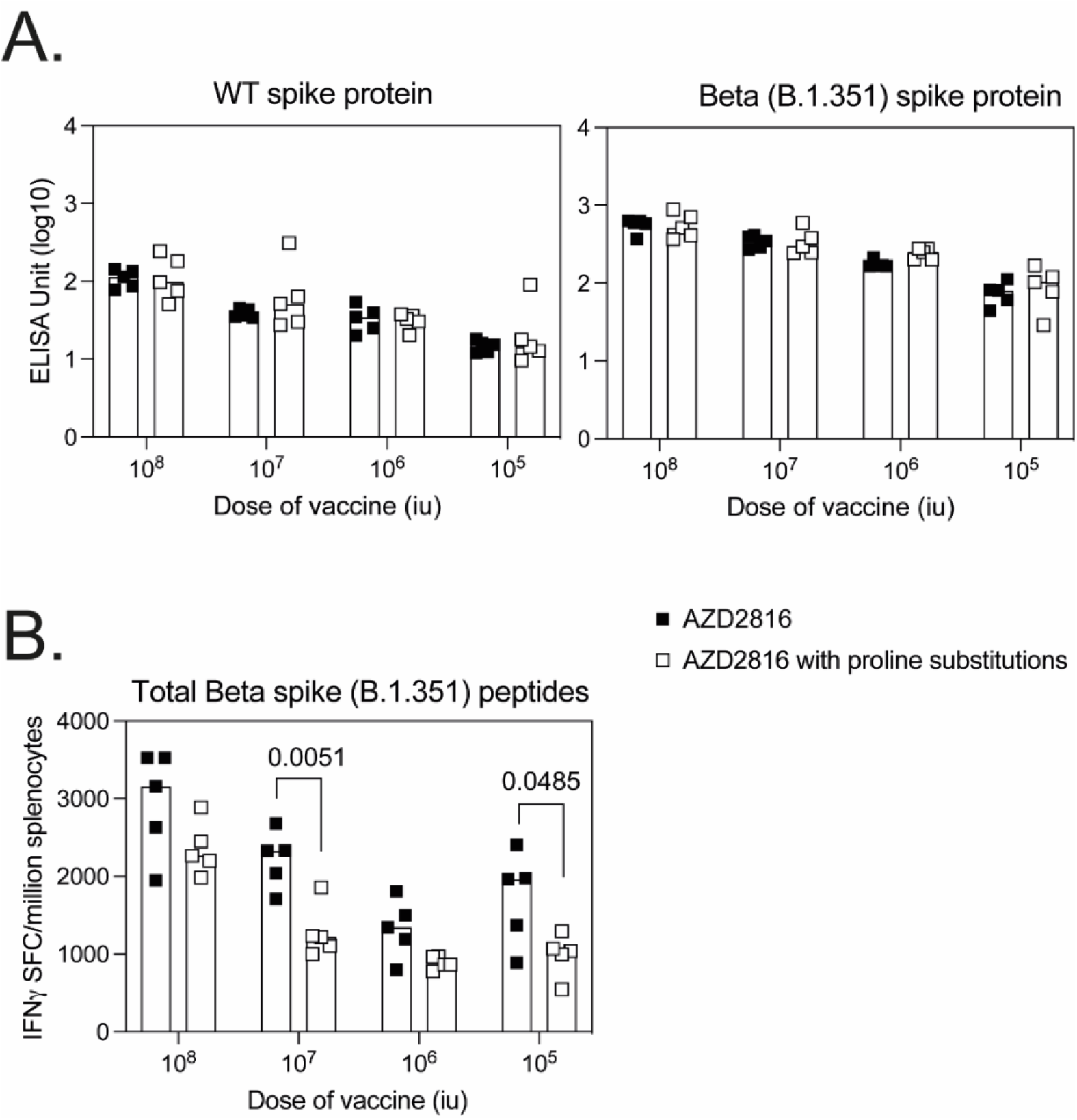
Comparison of ChAdOx1 expressing B.1.351 spike protein with or without proline substitutions. BALB/c mice were immunised with ChAdOx1 expressing SARS-CoV-2 B.1.351 spike protein (AZD2816) or B.1.351 spike protein containing 6 proline substitutions (AZD2816 without proline substitutions) across a range of doses. Mice were sacrificed 3 weeks later for measurement of antibody responses in the serum and cell-mediated responses in the spleen. **A**.) Graphs show total IgG antibody responses measured by ELISA against WT and B.1.351 spike protein, data was analysed with a two-way analysis of variance (repeated measure), no significant difference between vaccines was observed at any dose. **B**.) The graph shows the summed B.1.351 spike ELISpot response, data was analysed with a two-way anova and post-hoc positive test (Šidaks multiple comparison), p values denote statistically significant differences (p<0.05) between vaccines.

**Figure S2:**
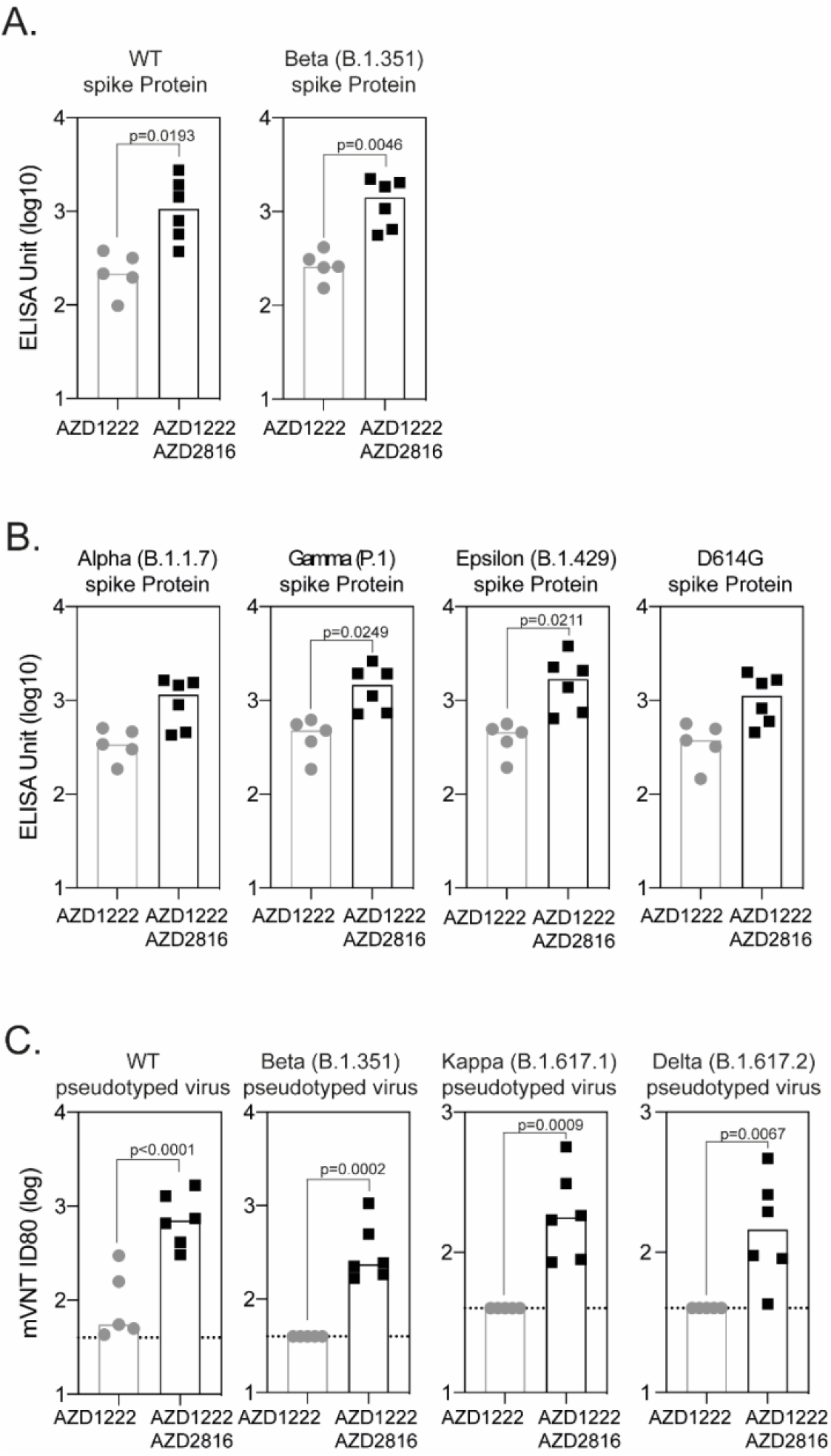
Antibody titres and breadth are increased following a booster dose with AZD2816 vaccine. **A**.) Graphs show total IgG response against original spike protein or Beta (B.1.351) measured in the serum of mice collected 16 days after vaccination with AZD1222 (n=5) (animals from Figure 2) or a prime-boost regimen of AZD1222 followed 4 weeks later by AZD2816 (n=6). **B**.) Graphs show total IgG responses measured against Alpha (B.1.1.7), Gamma (P.1), Epsilon (B.1.429) or D614G spike proteins in serum collected 16 days and 3 weeks after the final vaccination. All ELISAs were performed simultaneously, data log transformed and analysed with a 2-way anova with a post-hoc positive test, statistically significant differences between groups (p<0.05) are indicated. **C**.) Microneutralisation titre of serum (ID80) collected day 16 post-vaccination (animals Figure 2) and 21 days after prime-boost vaccination against pseudotyped virus expressing original, Beta (B.1.351), Kappa (B.1.617.1) or Delta (B.1.617.2) spike protein. Limit of detection in the assay is defined as a titre of 40 (dotted line). Data was log-transformed and analysed with a 2-way anova (repeated measure) and post-hoc positive test, statistically significant differences (p<0.05) between groups are indicated.

**Figure S3:**
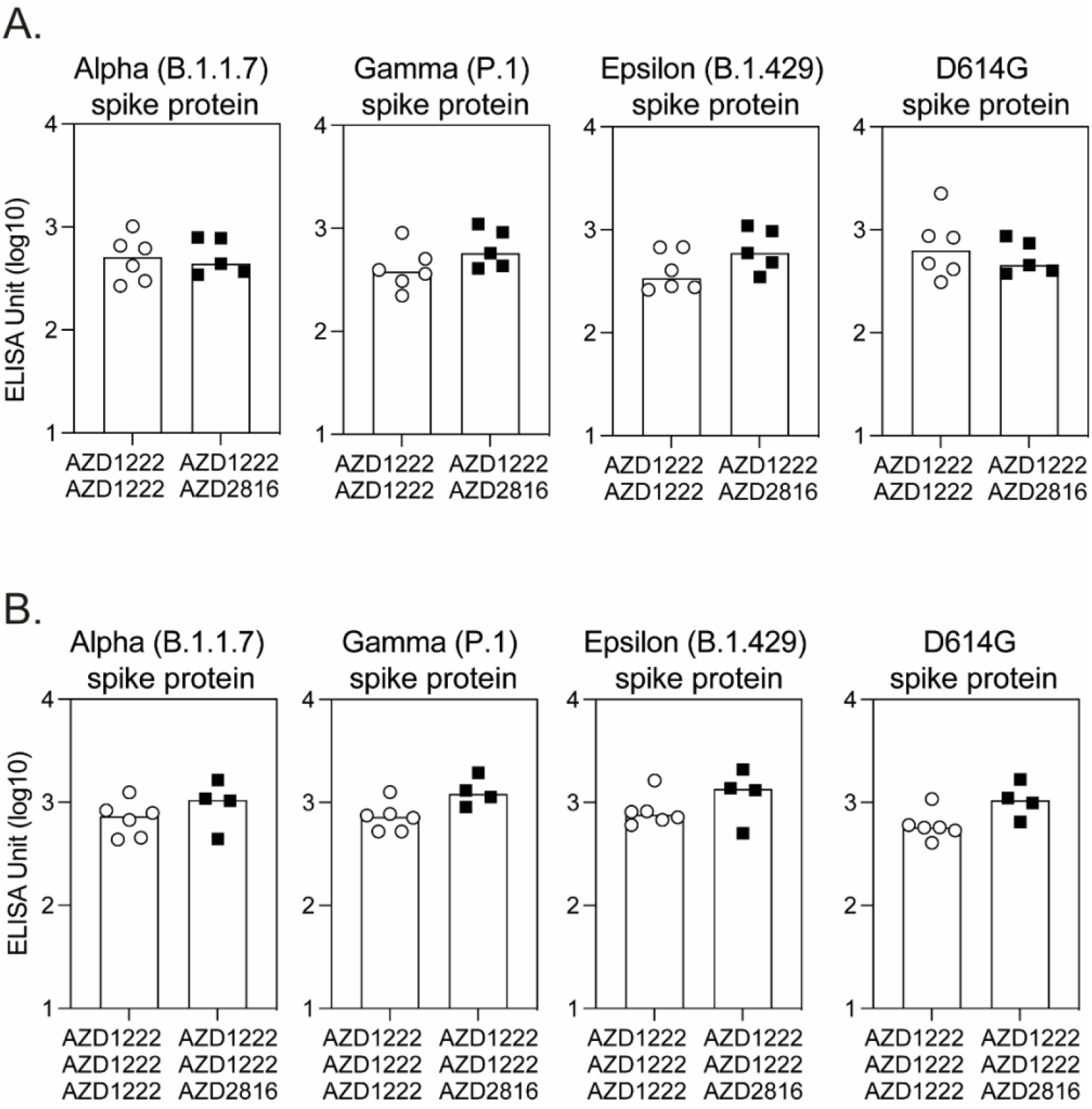
Breadth of antibody response to VoC proteins. **A**.) In the same experiment as described in Figure 3, BALB/c mice were primed with AZD122 and boosted 4 weeks later with either AZD1222 or AZD2816 and antibody responses to VoC protein measured by ELISA. Graphs show total IgG responses measured against Alpha (B.1.1.7), Gamma (P.1), Epsilon (B.1.429) or D614G spike proteins in serum collected 3 weeks after the final vaccination. **B**.) In the same experiment as described in Figure 4, BALB/c mice received two doses AZD122 prior to a final 3^rd^ dose of either AZD1222 or AZD2816, antibody responses to VoC protein were measured by ELISA 3 weeks after the final vaccination. Graphs show total IgG responses measured against Alpha (B.1.1.7), Gamma (P.1), Epsilon (B.1.429) or D614G spike proteins in serum collected 3 weeks after the final vaccination. All ELISAs were performed simultaneously, data log transformed and analysed with a 2-way anova to determine the effect of number of vaccine doses or booster vaccine. A statistically significant difference in ELISA Units observed between the number of doses of vaccines was observed against Alpha (p=0.0312), Gamma (p=0.0024), Epsilon (p=0.0042) but not D614G protein, no significant difference between booster vaccines was observed.

**Figure S4:**
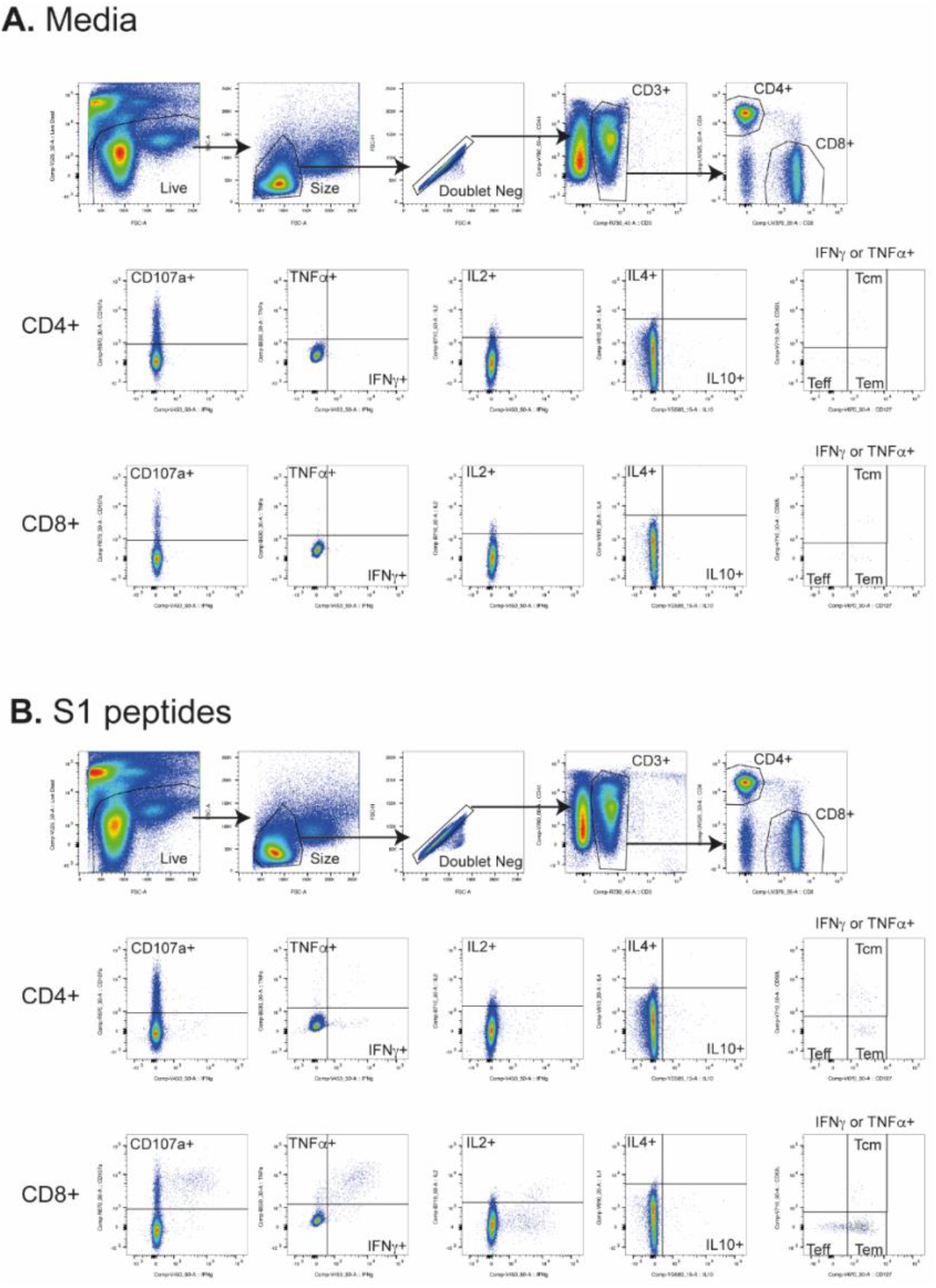
Flow cytometry gating strategy. Antigen specific T cells were identified by gating on LIVE/DEAD negative, size (FSC-A vs SSC), doublet negative (FSC-H vs FSC-A), CD3^+^, CD4^+^ or CD8^+^ cells and each individual cytokine. T effector (Teff) cells were defined as CD62L^low^ CD127^low^, T effector memory (Tem) cells defined as CD62L^low^ CD127^hi^ and T central memory (Tcm) cells defined as CD62L^hi^ CD127^hi^. T cell subsets were gated within the population of “IFNγ^+^ or TNFα^+^” responses and are presented after subtraction of the background response detected in the corresponding media stimulated control sample (**A**.) from the S1 (**B**.) or S2 peptide stimulated sample for each mouse and summing together the response detected to each pool of peptides.

